# The signal and the noise - characteristics of antisense RNA in complex microbial communities

**DOI:** 10.1101/776971

**Authors:** Thomas Yssing Michaelsen, Jakob Brandt, Caitlin Singleton, Rasmus Hansen Kirkegaard, Nicola Segata, Mads Albertsen

**Affiliations:** Center for Microbial Communities, Aalborg University; Department CIBIO, University of Trento

## Abstract

High-throughput sequencing has allowed unprecedented insight into the composition and function of complex microbial communities. With the onset of metatranscriptomics, it is now possible to interrogate the transcriptome of multiple organisms simultaneously to get an overview of the gene expression of the entire community. Studies have successfully used metatranscriptomics to identify and describe relationships between gene expression levels and community characteristics. However, metatranscriptomic datasets contain a rich suite of additional information which is just beginning to be explored. In this minireview we discuss the different computational strategies for handling antisense expression in metatranscriptomic samples and highlight their potentially detrimental effects on downstream analysis and interpretation. We also surveyed the antisense transcriptome of multiple genomes and metagenome-assembled genomes (MAGs) from five different datasets and found high variability in the level of antisense transcription for individual species which were consistent across samples. Importantly, we tested the hypothesis that antisense transcription is primarily the product of transcriptional noise and found mixed support, suggesting that the total observed antisense RNA in complex communities arises from a compounded effect of both random, biological and technical factors. Antisense transcription can provide a rich set of information, from technical details about data quality to novel insight into the biology of complex microbial communities.

**Key points:**

- Several fundamentally different approaches are used to handle antisense RNA
- Prevalence of antisense RNA is highly variable between communities, genomes, and genes.
- Antisense RNA is likely an opaque mixture of technical, biological and random effects

## Introduction

Transcriptomics is an established tool to quantify bacterial messenger RNA (mRNA) expression and regulation on a whole-organism level (Sorek and Cossart 2010), which increasingly also includes a wide repertoire of small regulatory non-coding RNA (sRNA) (Gorski, Vogel, and Doudna 2017; Hör, Gorski, and Vogel 2018). This is largely due to the introduction of next-generation RNA-sequencing (RNAseq), a technique initially applied mostly on human samples but which has also allowed unprecedented insight into the overall characteristics of complete microbial transcriptomes (Sorek and Cossart 2010). The probe-independent nature of RNAseq provides an unbiased means of identifying novel putative genes and sRNA elements which may be missing from the genomic annotation (Gorski, Vogel, and Doudna 2017). In particular, strand-specific RNAseq enables one to retain the directionality of the native RNA segment when sequenced and quantify antisense RNA (asRNA), here defined as the sRNA matching the template DNA strand inside a gene (Bao et al. 2015). Despite the wide array of studies identifying asRNAs as biological regulators of mRNA and transcription (Gorski, Vogel, and Doudna 2017), contrasting evidence suggests bulk asRNA is primarily the product of transcriptional noise, rising from spurious promotors (Lloréns-Rico et al. 2016). As bacterial promotors are characterized by low information content, these could arise by random mutations throughout the genome and cause transcription at constant rates across the genome without biological function (Lloréns-Rico et al. 2016; Wade & Grainger 2018).

Multiple studies have identified asRNA to be ubiquitous in bacteria (Thomason & Storz 2010; Sesto et al. 2013) and increasingly relevant in archaea (Gelsinger & Diruggiero 2018). These studies identified asRNA primarily through genome-wide searches for putative sRNA and transcriptome analysis of single-culture bacteria (Wagner & Romby 2015). Across different bacteria, the number of genes with asRNA varies substantially from more than 80% in *Pseudomonas aeruginosa* PA14 (Eckweiler & Häussler 2018) to 2.3% in Lactococcus lactis (van der Meulen et al. 2016). The functionality of asRNA has been extensively studied and several interactions between specific asRNAs and mRNAs have been described (Gorski et al. 2017). In Staphylococcus aureus it is suspected that at least 75% of mRNAs are modulated by asRNA (Lasa et al. 2011) and a substantial amount of asRNA-mRNA interactions were directly identified in *Escherichia coli* (Waters et al. 2016) and *Salmonella* (Förstner et al. 2016).

Pure culture studies are powerful tools to quantify asRNA and identify specific asRNA-mRNA interactions, but may not generalize to how the organism responds in complex microbial communities and its natural complex environment. The interplay between organisms changes their transcriptional pattern through response to e.g. competition or commensal interactions (Hibbing et al. 2010), but how this extends to asRNA remains largely unknown (Bashiardes et al. 2016). Detailed studies of the presence and dynamics of asRNA in complex microbial communities are scarce - to our current knowledge only one study investigated asRNA in human gut communities (Bao et al. 2015). Consequently, our understanding of the variability and persistence of asRNA in complex communities, between and within different environments, is limited.

The onset of metatranscriptomics enabled the transcriptome of multiple organisms to be examined simultaneously, providing a complete overview of the entire community gene expression without the need for culturing. This has made metatranscriptomics a powerful tool to detect and describe relationships between gene expression levels and community characteristics (Bashiardes et al. 2016). When coupled with metagenomics for the *de novo* construction of metagenome-assembled genome (MAGs) this provides a high-throughput culture-independent method of performing species-level analysis *in situ* (Mallick et al. 2017), not limited to genomes from already well-characterized species (Pasolli et al. 2019). As an increasing amount of metatranscriptomic data are being generated by strand-specific RNA sequencing, we encourage researchers to embrace this extra dimension of the data adequately. In this study we reviewed the different approaches for processing asRNA and highlight some of the emergent quantitative properties of asRNA, to provide a bird’s-eye view of antisense transcription in complex microbial communities.

## Methods

A total of 56 strand-specific metatranscriptomes from anaerobic digester (Hao et al. in preparation), water (Beier et al. 2017), permafrost soil (Woodcroft et al. 2018), and human gut (Bao et al. 2015) environments were considered and re-analyzed in this study. Metatranscriptomic reads were mapped to metagenome-assembled-genomes (MAGs) assembled from single samples or reference genomes provided by the respective studies (Table S1).

### Anaerobic digester

Samples originate from a lab-scale short-chain fatty-acid stimulation experiment. Slurry taken directly from an anaerobic digester at Fredericia wastewater treatment plant (55.552301, 9.721908) was incubated in reactors and subjected to a spike-in of acetate. Two samples for DNA sequencing on the Nanopore and Illumina platform was taken immediately before spike-in. Library preparation for Illumina sequencing is described elsewhere (Hao et al. in preparation). Library preparation for Nanopore sequencing was carried out using the SQK-LSK108 kit (Oxford Nanopore Technologies, UK) following the manufacturer’s protocol without the optional shearing step. The DNA libraries were sequenced using FLO-MIN106 MinlON flowcells (Oxford Nanopore Technologies, UK) on a MinlON mk1b (Oxford Nanopore Technologies, UK). The raw fast5 data was basecalled using Guppy v2.2.2 with flipflop settings (Oxford Nanopore Technologies, UK). The Nanopore reads were filtered removing short-reads (<4000 bp), then adapters were removed using Porechop vO.2.3 (Wick et al. 2017), and filtered based on quality using filtlong v0.1.1 (https://github.com/rrwick/Filtlong). The quality filtered reads were assembled using CANU v1.8 (Koren et al. 2017) circular contigs were aligned to themselves using nucmer v3.1 (Kurtz et al. 2004) and the overlap was trimmed manually. The assembly was polished with Nanopore reads using minimap2 v2.14 (H. Li 2018) and Racon v1.3.2 (Vaser et al. 2017) followed by polishing using medaka vO.4.3 twice (Oxford Nanopore Technologies, UK) and finally polished with minimap2 v2.14 and Racon vl.3.2 using Illumina data from (Hao et al. in preparation). The reads were then mapped back to the assembly using minimap2 v2.14 (H. Li 2018) to generate coverage depth files to assist automatic binning using metabat2 v2.12.1 (Kang et al. 2019). The generated MAGs were quality assessed using checkM v1.0.11 (Parks et al. 2015) with the ‘lineage_wf option. Gene detection and annotation were performed using prokka v1.13 (Seemann 2014) with standard settings apart from “--metagenome --kingdom Bacteria”. Sampling for RNA sequencing was done immediately before and at regular intervals up until 6 hours after spike-in. Experimental details and protocols for RNA sequencing are available in (Hao et al. in preparation).

### Water

Samples originate from a study by Beier and colleagues (Beier et al. 2017), who analyzed functional redundancy in fresh-, brackish-, and salt-water communities. DNA sequencing data was downloaded from the Sequence Read Archive (SRA) under accession number PRJEB14197. As some samples were sequenced multiple times, we only included the sequencing run with most reads for each sample. Sequence reads were trimmed for adaptors using cutadapt v1.16 (Martin 2011), and assembled using megahit v1.1.3 (D. Li et al. 2016). The reads were then mapped back to the assembly using minimap2 v2.12 (H. Li 2018) to generate coverage depth files to assist automatic binning using metabat2 v2.12.1 (Kang et al. 2019). A total of 147 MAGs were generated and quality assessed using checkM vl.0.11 (Parks et al. 2015) with the ’lineage_wf option. Gene detection and annotation were performed using prokka vl.13 (Seemann 2014) with standard settings apart from “--kingdom Bacteria --metagenome”. RNA sequencing data was downloaded from SRA under accession number PRJEB14197 and trimmed using cutadapt vl.16 (Martin 2011).

### Bog and Fen

Samples originate from a study by Woodcraft and colleagues (Woodcraft et al. 2018), who analyzed carbon processing in thawing permafrost soil communities. We choose to only analyse metatranscriptomes from bog and fen samples due to insufficient sample sizes in the other groups. We obtained the raw sense and antisense gene-expression data by personal communication with the authors, which were mapped and quantified as described in (Woodcraft et al. 2018). We also received the gene sequences for annotation. Assembly and quality statistics of all 1,529 MAGs generated in this study were available in Supplementary Data 3 from that study.

### Human gut

Samples originate from a study by Bao and colleagues (Bao et al. 2015), who analyzed antisense transcription in human gut communities. We downloaded sense and antisense gene-expression data from data sheet 2 in (Bao et al. 2015) available at http://journal.frontiersin.org/article/10.3389/fmicb.2015.00896. We focused only on the 47 species reported in Data Sheet 1 from (Bao et al. 2015), whose curated genomes were downloaded from an archived version of the FTP NCBI database (https://ftp.ncbi.nlm.nih.gov/genomes/archive/old_refsea/Bacteria). The genomes were quality assessed using checkM v1.0.11 (Parks et al. 2015) with the ‘lineage_wf option.

### Mapping and quantifying RNAseq data

trimmed RNA sequencing data from the anaerobic digester and water samples were mapped using bowtie2 v2.3.4.1 (Langmead and Salzberg 2013) to their respective metagenomes with settings ‘-a --very-sensitive --no-unal’. Downstream quantification was done using the python script BamToCount_SAS.py (https://github.com/TYMichaelsen/AntisenseStudv/blob/master/scriPts/BamToCount_SAS.py) parsing the bam files using pysam v0.15.0 (https://github.com/pysam-developers/pysam) and gff files using Biopython v1.72 (Cock et al. 2009).

### Filtering and preprocessing expression data

After acquisition of data from all four studies we retained only samples with a total read count of >1 million reads. Each dataset was then filtered separately by the following steps: 1) include only protein coding genes; 2) remove genes with zero counts in >75% of samples to filter out rarely expressed genes; and 3) remove observations with total read count < 10 to filter low-expressed genes with low signal-noise ratio. Transcripts per million (TPM; (Wagner, Kin, and Lynch 2012)) were calculated from these datasets. Next, a binomial test for significant antisense transcription was done as described in (Bao et al. 2015). Specifically, artifacts introduced by cDNA synthesis and amplification are known problems for antisense RNA detection (Thomason and Storz 2010), which leads to false positive detection of antisense transcription. Let *q* be the probability of having antisense reads when in reality there is none (i.e. the inherent technical error/false positive rate). We then have a null hypothesis of observing *q* or less (i.e. a one-tailed test) and test if the true percentage of antisense reads deviates from this. If the observed p-value, after correction using the Benjamini-Hochberg procedure (Benjamini and Hochberg 1995), is less than 0.05 we consider that the gene has antisense transcription. We used a *q* value of 0.01 as suggested by (Bao et al. 2015), but tested also *q* = 0.05 as a more conservative estimate for comparison. From here, two datasets were generated for each community - one fulfilling previous criteria (sample-wide dataset) and another specifically to analyse genome-wide antisense expression (genome-wide dataset). The genome-wide dataset was subjected to an additional filtering. First, all genomes/MAGs failing to reach the criteria for a medium-quality MAG (completeness ≥ 50% and contamination < 10%) determined by the Genomic Standards Consortium (Bowers et al., n.d.) were removed. Second, all genomes with <50 genes above detection limit (total read count ≥ 10) were removed for each sample separately. This was done to ensure enough genes were above the detection limit to accurately calculate genome-wide estimates. Furthermore, all genes referenced by name in this manuscript were manually validated against the NCBI database (https://www.ncbi.nlm.nih.gov/nucleotide) to support correct annotation.

### Enrichment analysis

We used the Cluster of Orthologous Groups (COG) database (Galperin et al. 2015) as the basis for enrichment analysis. The full database was downloaded from the NCBI FTP archive at ftp://ftp.ncbi.nih.gov/pub/COG/COG2014/data. To test for enrichment/reduction of a specific COG category in a set of genes *n*, drawn from the entire population of genes *N*, a hypergeometric test was performed. We calculated the probability of having k genes associated to the given COG in the set *n*, given that there were *K* genes in the remaining population *N - n*. If the p-value for a one-tailed test for over- or under-representation of the COG category was below 0.05 after Benjamini-Hochberg correction (Benjamini & Hochberg 2019) we considered it to be enriched or reduced, respectively. Two-tailed tests were performed by calculating the p-values for both tails, taking the smallest value and multiplying by two (Rivals et al. 2007).

## Results and discussion

### Three approaches to handle asRNA in literature

We identified three common approaches to handle strand-specific metatranscriptomic RNAseq data (Table 1), which rely on substantially different assumptions about the origin of asRNA and its effect on study outcome. First, some studies ignore the strand-specific information of the data completely (Franzosa et al. 2014, 2018; Asnicar et al. 2017; Beier et al. 2017), implying that the expression of a given region will be the sum of the sense and antisense expression. This simplifies the downstream analysis and has been the traditional approach given that stranded RNAseq, conserving the sense of RNA transcripts, is relatively new (Levin et al. 2010). Importantly, it is also based on the assumption that asRNA is irrelevant for the study outcome, intentionally or not. Second, other studies filter away the antisense mapping reads (Heintz-Buschart et al. 2017), which is also integrated into many popular tools designed for quantifying gene expression such as HTSeq (Anders, Pyl, and Huber 2015), Cufflinks (Trapnell et al. 2010), and featureCounts (Liao, Smyth, and Shi 2014). This ensures quantification of mRNA only. Third and finally, there are studies which directly incorporate the strand-specific signal into the analysis using statistical tests comparing the sense and antisense expression within each region of interest (Bao et al. 2015; Woodcroft et al. 2018). This may be used to filter away genes with low sense relative to antisense expression (Woodcroft et al. 2018), or to identify genes with significant antisense expression (Bao et al. 2015).

**Table 1.** Listing of relevant papers and their procedure to handle antisense RNA.

| Study | Environment | Strandness | Handling |
| --- | --- | --- | --- |
| (Woodcroft et al. 2018) | Soil | Scriptseq Complete kit (bacteria) | Incorporated |
| (Bao et al. 2015) | Human gut | Custom protocol* | Incorporated |
| (Heintz-Buschart et al. 2016) | Human gut | NEBNext Ultra Directional RNA kit | Filtered |
| (Beier et al. 2017) | Water | TruSeq Stranded kit | Ignored |
| (Franzosa et al. 2018) | Human gut | RNAtag-seq** | Ignored |
| (Asnicar et al. 2017) | Human gut | ScriptSeq Complete Gold kit | Ignored |
| (Angle et al. 2017) | Soil | Truseq Stranded kit | Ignored |
| (Franzosa et al. 2014) | Human gut | RNAtag-seq** | Ignored |
| * See (Giannoukos et al. 2012) |  |  |  |
| ** See (Shishkin et al. 2015) |  |  |  |

### Antisense RNA is concentrated on few genes which are functionally reduced

To elucidate asRNA in strand-specific metatranscriptomes, we did an explorative meta-analysis of the quantitative, functional and genome-specific properties of asRNA in five public datasets covering an anaerobic digester, water, bog and fen permafrost-associated wetlands, and human gut. The percentages of asRNA in the datasets varied from 1.1% to 10.4% for the human gut and water communities (Figure 1A). The anaerobic digester, bog, and fen samples had a substantially higher percentage of asRNA ranging from 33.2-90.8%. Surprisingly, only 2-3 genes accounted for up to 90% of all asRNA in these communities (Figure 1B). Among three genes with the highest median percentage of asRNA across all communities, 3 of them could be annotated as cytadherence high molecular weight protein 2 (hmw2; max-min range: 59% - 61%), a putative protein (max-min range: 17.5% −18%), and M protein, serotype 5 (emm5; max-min range: 11% - 11.5%). Both hmw2 and emm5 can act as virulence factors (St Gerne and Yeo 2009; Luca-Harari et al. 2009). Conversely the abundance of the three genes with highest median mRNA expression relative to total mRNA were highly variable (anaerobic digester: 6% - 21%; bog: 14% - 57%; fen: 8% - 57%). We found no spurious amplifications of primer regions when inspecting the coverage profiles of these genes, making it unclear what causes the consistent high expression of asRNA. Generally, the cumulative percentages of mRNA follows a log-linear dependence on the number of genes for all five communities (Figure 1B). By removing the 10 most asRNA-expressing genes from analysis (Figure S1) the percentage of asRNA was drastically reduced in the anaerobic digester, bog and fen samples and the cumulative percentage profiles approached those of human gut and water samples.

**Figure 1.**
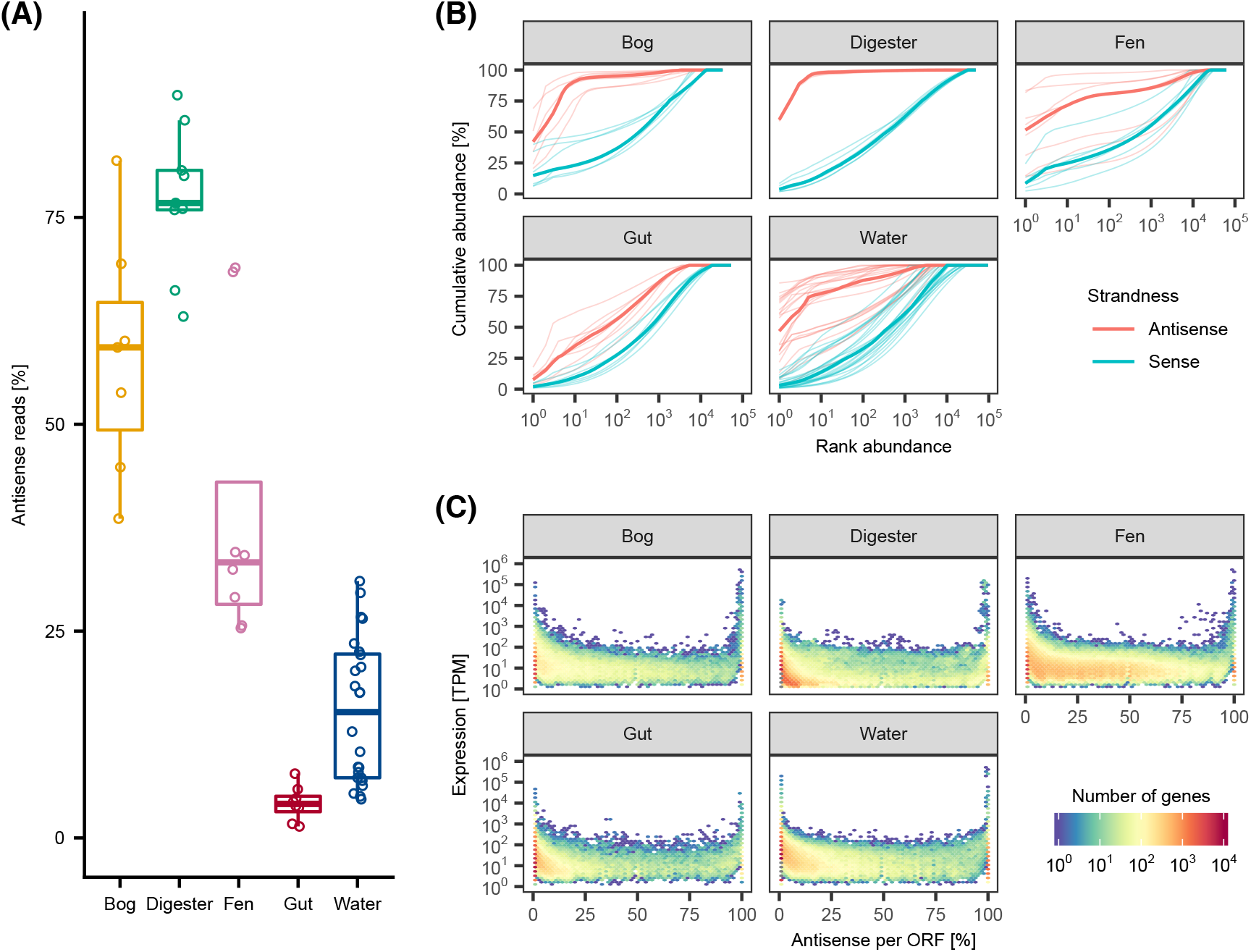
Antisense transcription is a substantial amount of the data and dominated by few genes. **(A)** Percentage of sequenced reads which are antisense for each sample, grouped by community. (B) Rank-abundance curves for all samples within each community, colored by strandness. The x-axis lists genes ranked in descending order according to relative abundance, while the y-axis shows the cumulative abundance. Bold lines indicate the median for each colored group. (C) On the x-axis the relative amount of antisense expression is shown for each observation while the y-axis shows the total antisense and sense expression normalized to transcripts per million (TPM). For this figure, the sample-wide datasets were used (see Methods for detail).

By plotting the combined mRNA and asRNA expression against the percentage of asRNA for each gene we observed a characteristic U shape across all five communities (Figure 1C). A substantial percentage of gene expression mRNA/asRNA pairs are exclusively mRNA or asRNA in the anaerobic digester (mRNA only = 73.8%, asRNA only =1.4%), bog (mRNA only = 60.5%, asRNA only = 6.3%), fen (mRNA only = 26.3%, asRNA only = 4.2%), human gut (RNA only = 73.4%, asRNA only = 1.2%), and water (RNA only = 79.7%, asRNA only = 2.7%) communities. Generally, genes with either pure RNA or asRNA expression tend to be highly expressed, which has been reported previously (Bao et al. 2015). We now show this to be a consistent phenomenon across different environments, suggesting biological relevance. The presence of asRNA might have regulatory properties, as high levels of asRNA could act to block gene expression (Bao et al. 2015; Hör, Gorski, and Vogel 2018). Such information could for example partition genes into suppressed, intermediate, and unsuppressed based on certain cutoff values of percent asRNA (Bao et al. 2015). Alternatively, the high percentage of genes with asRNA in bog and particularly fen samples have previously been attributed to DNA contamination, as double-stranded DNA synthesized from mRNA during library preparation may separate, causing information of strandness to be lost (Woodcroft et al. 2018). This approach discards large proportions of the data upfront and may be too conservative for many purposes. Furthermore, DNA contamination due to technical errors should occur randomly (Thomason and Storz 2010) and therefore evenly for all genes cancelling out any biases. However, no study to our knowledge has assessed the extent of DNA contamination due to technical artifacts in complex metatranscriptomes.

To identify potential high-level biological drivers of asRNA expression we investigated enrichment of COG functional categories in the five different environments. Antisense-enriched genes, which were defined as genes with asRNA accounting for >95% of the total RNA, were compared against the remaining background of genes (Figure 2A). All COG categories except cytoskeleton (category Z) and nuclear structure (category Y) were detected in all communities (Figure 2A). Between 23% - 48% (Bog: 23%; Anaerobic digester: 43%; Fen: 24%; Human gut: 29%; Water: 48%) of all genes with expression above detection limit were assigned COG identifiers. The majority of COG categories were significantly reduced in asRNA-enriched genes across all five communities, while the prevalence of genes without COG annotation were significantly enriched (all communities: p < 0.001) from a mean of 67% (52% - 77%) to 87% (85% - 90%) (Figure 2A). This contrasts the findings of Bao and colleagues (Bao et al. 2015), who identified multiple COG categories to be enriched. However, direct comparison with the results of this study is not possible as we used the ratio of asRNA directly opposed to a statistical test of significant asRNA expression used by (Bao et al. 2015). Regardless, this lack of functional assignment demands further research. Linking levels of asRNA to the suppression/activation of certain genes or pathways in a case-control setup might provide a path forward to integrate such information into metatranscriptomic studies (Bashiardes, Zilberman-Schapira, and Elinav 2016).

**Figure 2.**
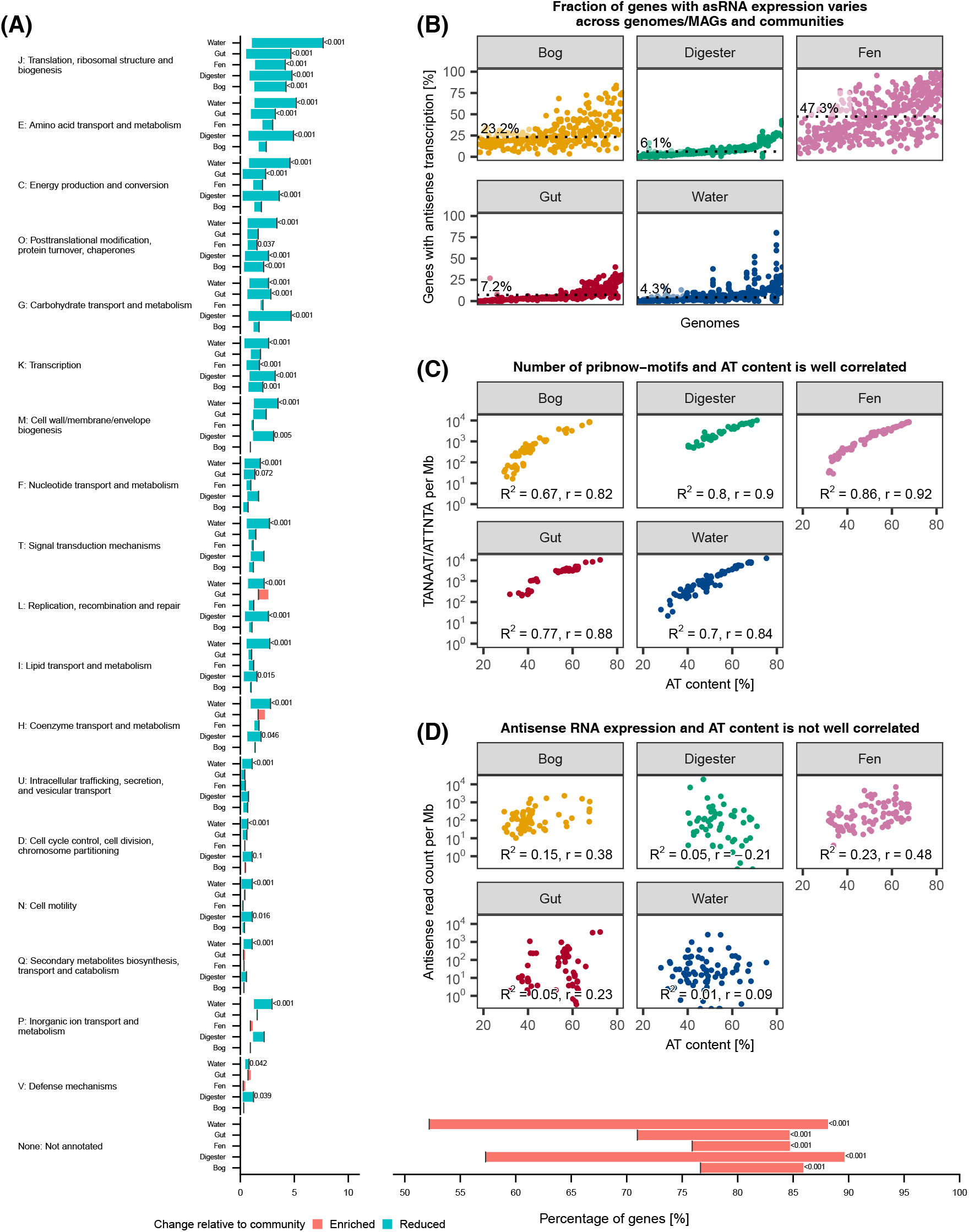
Antisense transcription is not function specific or due to transcriptional noise. (A) Listed from top to bottom is the enrichment/reduction for each COG functional category for each community, while the x-axis shows the percentage of genes in either the COG functional category or the entire community. Note the break between 10% and 50%. The black vertical line at each horizontal bar indicates the percentage of annotated genes in the whole community, while bars indicate either percent reduction (blue) or enrichment (red) in the subset of antisense-enriched genes, defined as genes with >95% reads being antisense RNA. Only two-tailed adjusted p-values < 0.2 are shown (see methods for details). Total number of genes and antisense-enriched subset in Digester [total: 48889, subset: 936 (1.9%)], Human gut [total: 53059, subset: 927 (1.8%)], Water [total: 95695, subset: 5339 (5.6%)], Fen [total: 62381, subset: 1759 (2.8%)], and Bog [total: 33258, subset: 1470 (4.4%)]. (B) For each community, genomes are plotted along the x-axis and the percentage of genes with significant (adjusted p-value < 0.05) antisense expression within each genome plotted on the y-axis. Points located above the same genome represent the same genome across different samples. For each community, genomes are ordered along the x-axis according to the median. Dotted lines and values indicate the median genome-wide percentage of genes with antisense transcription for each community. (C-D) On the x-axis are the percent AT-content for a given genome, while the y-axis shows for (C) the number of Pribnow-motifs in both directions of the sequence, normalized by genome length. (D) Shows the number of antisense RNA read counts normalized by genome length. Each dot represents a genome, whose value is the median across samples. For this plot the genome-wide datasets were used, see Methods for details.

### Levels of asRNA are genome-specific and not driven by transcriptional noise

We also investigated the variability in antisense expressing genes between genomes and samples. To this end, we quantified the percentage of genes with significant asRNA expression (adjusted p-value < 0.01) using a binomial test of the sense and antisense read count for each gene in each genome/MAG (Figure 2B). This varied substantially between communities, with an average from 4.3% to 47.3%. We used a two-way ANOVA within each community to partition the variance in genome-wide asRNA expression into sample and genome variability. We found that the variance explained by genome was greater than what could be attributed to sample variability in all five communities (anaerobic digester: genome = 88.3%, sample = 0.9%; bog: genome = 52.7%, sample = 26.2%; fen: genome = 45.3%, sample = 40.1%; human gut: genome = 63%, sample = 11%; water: genome = 43%, sample = 8.5%). The percentage of genes with significant asRNA expression and results of the ANOVA were consistent using a less conservative adjusted p-value of < 0.05 (Figure S2).

Genome-specific asRNA levels have been reported for pure cultures (Lloréns-Rico et al. 2016) and our study is the first to extend these findings beyond the human gut community (Bao et al. 2015). Thus, high inter-community variability of genome-wide asRNA expression is a prevalent phenomenon. Such differences may have a profound effect when comparing gene expression levels between genomes and can bias the analysis if not properly accounted for. Interestingly, the extent of this may also be community-specific, as genomes explained 43% to 88% of the variability in the percentage of genes with significant asRNA expression in water and anaerobic digester communities, respectively.

We attempted to extend the results of (Lloréns-Rico et al. 2016) to complex communities, stating that bulk asRNA is primarily the product of transcriptional noise, rising from spurious promoters. We found a log-linear dependency of the number of Pribnow motifs, normalized by genome size on the AT content in all communities (Figure 2C). This supports the hypothesis that an increase in AT content leads to an increase in spurious promoter sites. However, we did not find support for the positive effect of AT content on the expression of asRNAs normalized by genome size (Figure 2D), and the data does not support an exponential model as provided by (Lloréns et al. 2016) nor a simple linear model. Furthermore, we observed large variation in the asRNA levels across the AT content gradient (Figure 2D). This high variability might arise as a combined effect of multiple factors influencing the asRNA transcription in addition to pervasive transcription, emphasizing the complexity of metatranscriptomic data.

### The role of asRNA in metatranscriptomics research

It is not clear to what extent asRNA in metatranscriptomics reflects a biological rather than a stochastic process (Lloréns-Rico et al. 2016). Although a considerable body of literature is increasingly explaining the potential biological mechanisms behind asRNA, one should be careful generalizing these findings to metatranscriptomics. Reference genomes are often not available, meaning that genomes are constructed de novo from corresponding metagenomes (Albertsen et al. 2013). This often leads to incomplete annotation due to novelty, partially hampering downstream biological interpretation. Much of the current knowledge about asRNA is indeed based on pure cultures (Bashiardes, Zilberman-Schapira, and Elinav 2016), which may not be representative of the intricate interactions that potentially occur in coexistence with other organisms in complex communities, where single cells within a population may be in different states of growth and dormancy simultaneously (Shank 2018). Regardless, profiling microbial communities by high-throughput omics techniques remains a powerful tool to build metabolic networks and infer interactions between different organisms (Mallick et al. 2017; Franzosa et al. 2018; Muller et al. 2018) and we believe asRNA expression should be integrated into such analysis. From our literature search we found three approaches to handle asRNA (Table 1), of which ignoring asRNA or quantifying only the sense RNA are by far the most common. Ignoring strand-specificity might be reasonable for the majority of genes, given that these do not express asRNA (Figure 1C). However, metatranscriptomic studies are often explorative and as genes with mainly asRNA tends to be highly expressed, these are likely to be emphasized during standard genome-wide association studies (GWAS) and constitute a large proportion of the data. Such genes would be false positives or misinterpreted as being expressed in the traditional way (Bao et al. 2015). Quantifying only the directly transcribed RNA avoids such false positives and should be the naive, bare minimum requirement for quantifying gene expression in metatranscriptomes. Of course, we believe the optimum would be to include asRNA as a separate entity into the analysis whenever possible, as important Information may be retrieved. For example, high levels of asRNA could have inhibitive effects on downstream protein production (Gorski, Vogel, and Doudna 2017; Hör, Gorski, and Vogel 2018). This could be investigated by including asRNA in standard differential expression analysis and screening for significant interactions between asRNA/mRNA pairs and their association with experimental treatment.

### Future guidelines

It is of great interest to determine the biological or technical processes driving asRNA and RNA relationships in metatranscriptomic data (Wade and Grainger 2014; Lloréns-Rico et al. 2016; Hör, Gorski, and Vogel 2018). In this study, we reviewed and summarized the characteristics of asRNA in metatranscriptomes from complex microbial communities. We have provided simple statistics of prevalence and quantities, and elucidated the potential role and function of asRNA. This study provides a methodological foundation for addressing asRNA in metatranscriptome analysis in general. As we have shown, asRNA expression does not seem to be caused by spurious promoters and may vary substantially between communities on multiple parameters such as functional categories, sequence diversity and prevalence. If this variability is not acknowledged then important causal relationships could be overlooked and incorrect interpretation of data could be performed. For example, transcription of asRNA may constitute a significant percentage of the data and be associated to only a few genes. As sequencing data is inherently compositional, there will be an overrepresentation of spurious negative correlations with the remaining gene population, which cannot be amended using traditional quantitative data analysis (Quinn et al. 2018). This is true regardless of whether the highly expressed genes are systematically related to the experiment or not. Such issues will be avoided by quantifying mRNA only, ensuring there is sufficient data for proper mRNA quantification. However, compositional data analysis methods do exists, for which these issues can be amended (Erb et al. 2017; Quinn et al. 2018).

This study highlights that very little is known about asRNA expression in complex communities. Hypothesis-driven studies are warranted to investigate the function and characteristics of asRNAs in metatranscriptomes. Integrating asRNA analysis with existing methodologies and pipelines, such as GWAS and differential expression analysis, will facilitate clarification of its role in cell processes and regulation (Sesto et al. 2013). We believe asRNA in complex communities is an important part of the metatranscriptome machinery and interesting discoveries may lie ahead, driven by the approaches highlighted in this study.

## Acknowledgements

Thanks to Joel A. Boyd for providing the raw gene-expression and annotation data from the study by Woodcroft and colleagues (Woodcraft et al. 2018), as well as personal communication about study details. Thanks to Erika Yashiro for performing the Illumina sequencing of anaerobic digester samples.

---

**Figure S1.**
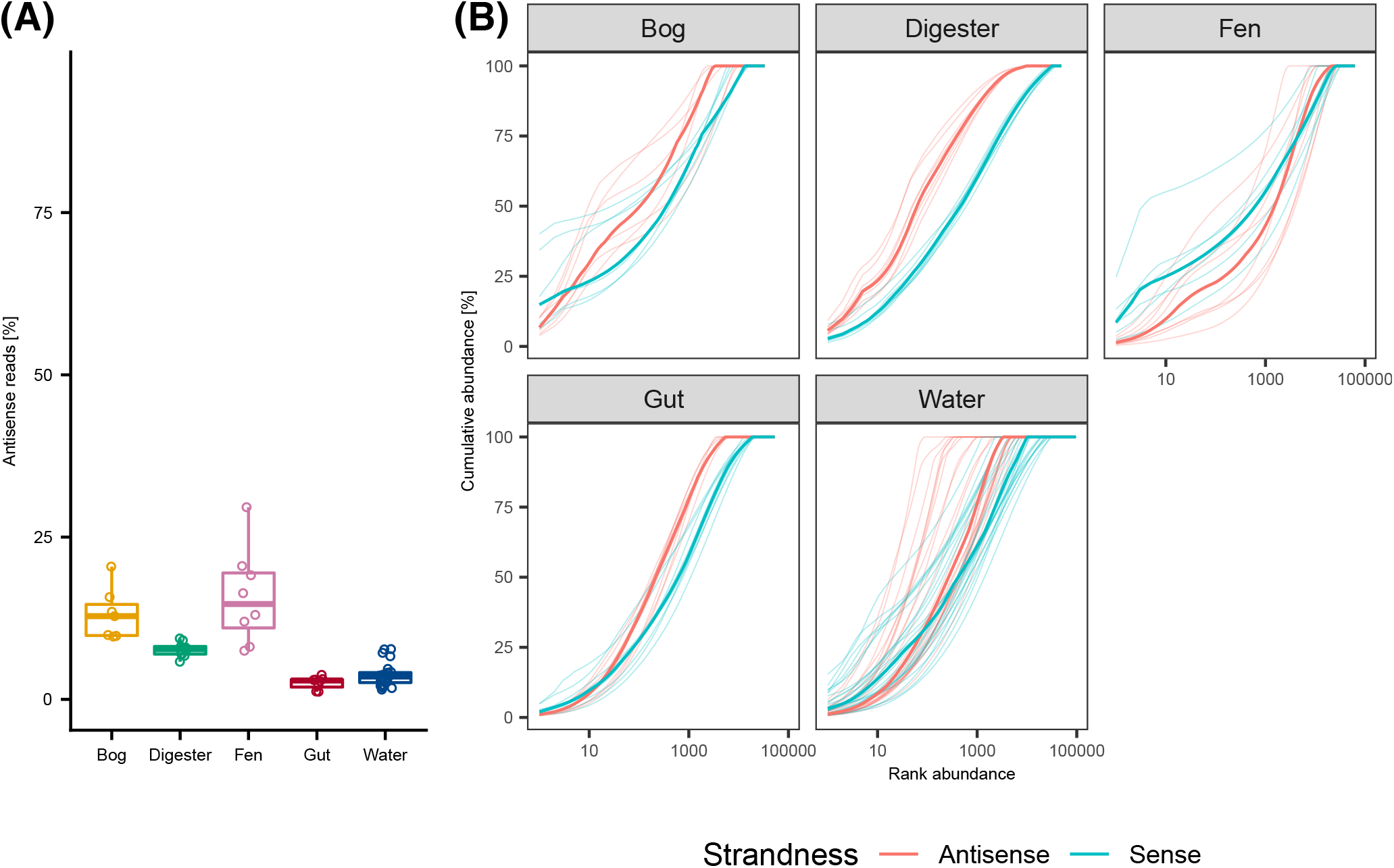
Antisense transcription is a substantial amount of the data and dominated by few genes (top 10 genes removed). (A) Percentage of sequenced reads which are antisense for each sample, grouped by community. (B) Rank-abundance curves for all samples within each community, colored by strandness. The x-axis lists genes ranked in descending order according to relative abundance, while the y-axis shows the cumulative abundance. Bold lines indicate the median for each colored group. For this figure, the sample-wide datasets were used (see Methods for detail).

**Figure S2.**
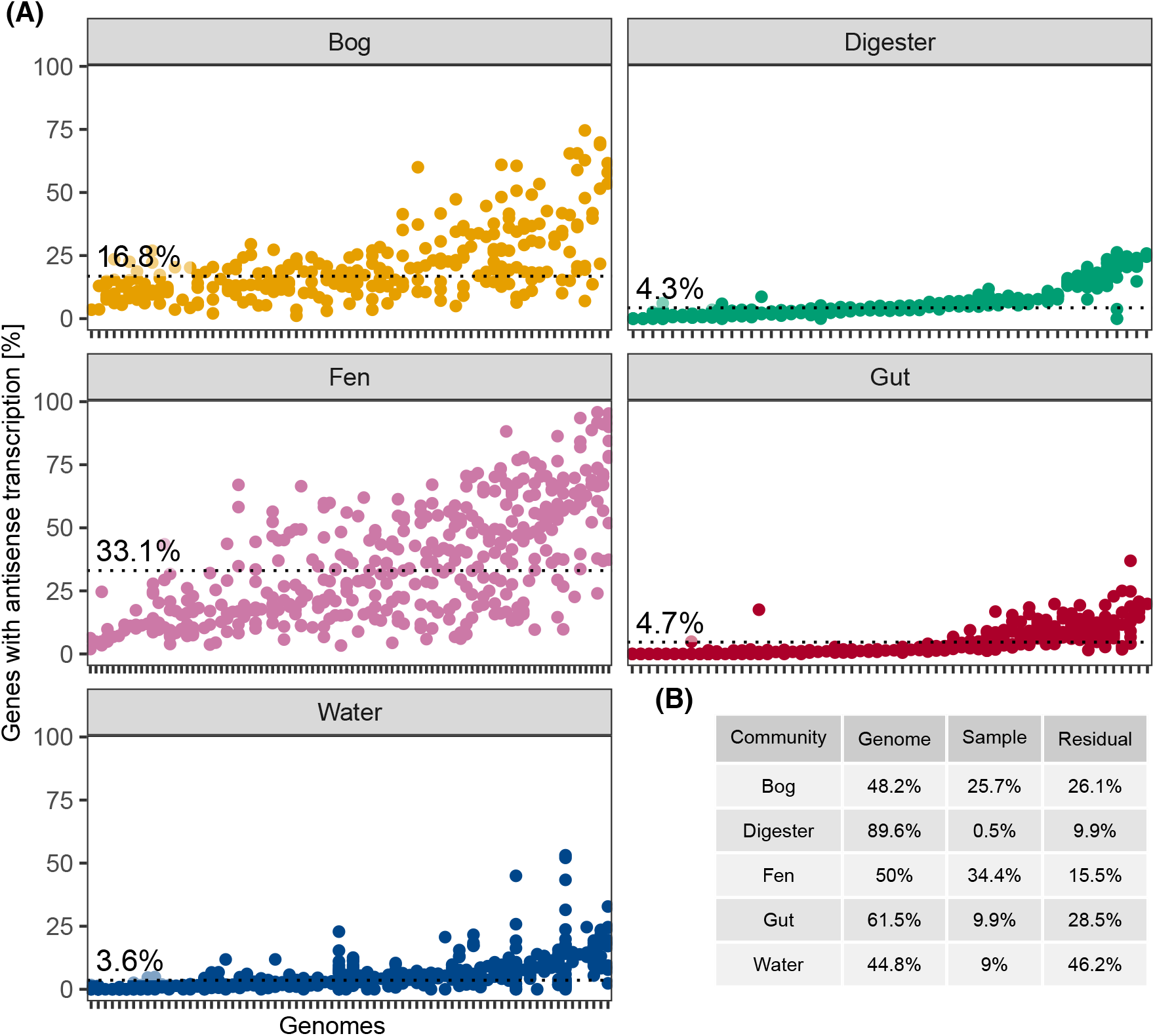
Genome-wide antisense transcription varies across and within samples (q-value = 0.05). (A) For each community, genomes are plotted along the x-axis and the percentage of genes with significant (adjusted p-value < 0.05) antisense expression within each genome on the y-axis. Points located above the same genome are representing the same genome across different samples. For each community, genomes are ordered along the x-axis according to the median. Dotted lines and values indicate the median genome-wide percentage of genes with antisense transcription for each community. (B) Explained variance output from a two-way ANOVA of genome and sample, performed on each community separately using the data in (A). For this plot the genome-wide datasets were used, see Methods for details.

**Table S1.** Characteristics of MAGs from the five different environments.

|  | **Digester** | **Bog** | **Human gut** | **Water** | **Fen** |
| --- | --- | --- | --- | --- | --- |
| **Samples** | 9 | 7 | 8 | 24 | 8 |
| **Genomes/MAGs** | 51 | 69 | 62 | 74 | 92 |
| **AT content (%)** | 53 (40 - 71) | 40 (29 - 68) | 57 (32 - 72) | 49 (28 - 75) | 52 (32 - 68) |
| **Size (Mb)** | 2.3 (1 - 3.9) | 4.3 (1.5 - 8) | 3.3 (1.8 - 6.3) | 2.9 (0.7 - 7.7) | 3.2 (1.5 - 8.7) |
| **Genes** | 2089 (1006 - 3394) | 3868 (1622 - 7176) | 2830 (1591 - 4902) | 2729 (768 - 6573) | 3039 (1622 - 8069) |
| **Completeness (%)** | 88 (57 - 98) | 95 (71 - 100) | 99 (76 - 100) | 89 (51 - 99) | 90 (71 - 100) |
| **Contamination (%)** | 1 (0 - 10) | 2 (0 - 9) | 0 (0 - 3) | 1 (0 - 10) | 2 (0 - 9) |

